# DPCMHC: efficient prediction of MHC-peptide binding affinity by deep learning based on dual-padding convolution

**DOI:** 10.64898/2025.12.01.691752

**Authors:** Yi Lu, Le Mi, Shixiong Zhang

**Affiliations:** School of Computer Science and Technology, Xidian University, Xi’an, Shaanxi, China; Department of Pharmacy, Xinle City Hospital, Xinle, Hebei, China

**Keywords:** MHC, peptides, deep learning

## Abstract

Predicting peptide binding affinities to major histocompatibility complex class I (MHCI) is a crucial challenge in immunological bioinformatics and is essential for identifying neoantigens in personalized cancer vaccines. Current deep learning methods often struggle, especially with 10-mer and 11-mer peptides. To address this, we developed DPCMHC, an advanced deep learning model featuring an embedding module, a dual-padding convolutional module, a BiLSTM module, and an output module. This design enhances the model’s understanding of amino acid sequences and their lengths. DPCMHC effectively captures the information on the beginning and end of amino acids, as well as the diverse sizes of adjacent amino acids. Using concatenation, the model extracts continuous sequence information. We rigorously evaluated DPCMHC on three benchmark datasets, demonstrating its superior or comparable performance to existing state-of-the-art methods. Our validation results confirm that DPCMHC is a robust and efficient tool for predicting MHC-peptide binding affinities.

## INTRODUCTION

The major histocompatibility complex (MHC) is crucial in the adaptive immune system ^1^, particularly in antigen presentation to T cells. Within the endoplasmic reticulum, peptides undergo trimming to fit specific MHC molecules. The peptide-MHC complexes (pMHC) are then assembled with the help of various chaperones and accessory proteins, ensuring correct folding and stability. Those complexes are transported to the cell surface. This process is vital for T cell activation and the targeting of infected cells, which is central to cellular immunity. Therefore, accurately predicting the affinity between MHC molecules and peptides is essential for developing vaccines, personalized medicine, and immune therapies ^2^.

MHC molecules are categorized into MHC class I (MHCI) and MHC class II (MHCII) ^3^. MHCI molecules primarily bind and present shorter peptides, typically 8-15 amino acids long, derived from endogenous proteins to CD8+ T cells ^4^. In contrast, MHCII molecules bind and present longer peptides, usually 13-25 amino acids ^5^, processed from exogenous proteins and recognized by CD4+ T cells ^6 7 8^. This difference in peptide length influences their respective roles in immune responses against different pathogens.

In the past years, many prediction algorithms have been developed for MHC-peptide binding prediction with many successful results ^9 10 11^. Most existing computational methods for predicting MHCI and peptide binding can be divided into two categories: pan-specific and allele-specific ^12^. Allele-specific methods have limited generalization for unseen MHCI molecules because they use separate models for each molecule ^13 14^. In contrast, pan-specific methods integrate information about both MHC molecules and peptides into a single model, allowing simultaneous learning of binding specificities across all MHC molecules ^15^. Those methods typically use binding assay data from the Immune Epitope Database and Analysis Resource (IEDB) ^16^, which has a significant bias toward peptides of length 9^10^. Many algorithms are designed specifically for 9-mer peptides and perform poorly with other lengths. Despite advances in machine learning, including methods like DeepAttentionPan ^17^, CaspNet-MHC ^18^, DeepMHCI ^19^, and DeepMHCI, challenges remain in predicting binding for non-9-mer peptides. Among these, DeepMHCI has shown the best performance for both 9-mer and non-9-mer.

Here we introduce DPCMHC, a deep-learning-based method for predicting the binding affinity of MHCI and peptide by integrating biological knowledge into model design. DPCMHC focuses on three key aspects: (i) dual-padding before convolution, (ii) concatenating after convolution, and (iii) different kernel sizes. Through dual-padding, the receptive fields were different for the two branches of one convolutional operation, which makes the model consider not only the beginning and ending amino acid information but also the adjacent amino acids of varying sizes. Additionally, with the concatenation operation, the model could better extract continuous sequence information after convolution.

We evaluated DPCMHC using three benchmark datasets and compared it with three state-of-the-art pan-specific methods. Our results showed that DPCMHC outperformed those methods in most experiments, especially on BD2017 for 9-mer, 10-mer, and 11-mer peptides. Furthermore, DPCMHC demonstrated superior performance and interpretability in identifying binding motifs of MHCI molecules across different peptide lengths.

## METHODS AND MATERIALS

### DATASETS

Three publicly available benchmark datasets are used to train and evaluate DPCMHC and benchmark methods. BD2017 served for model training and analyzing MHC-peptide binding affinities. Binary_2024_ acted as an independent dataset for assessing and classifying MHC-peptide binding affinities, while IEDB2016 evaluated the universal applicability of the DPCMHC architecture. These datasets collectively provide a robust foundation for evaluating DPCMHC and demonstrate its effectiveness across different peptide lengths and MHCI molecules.

#### BD2017

BD2017 was compiled from IEDB ^16^ and contains 185,985 peptide-MHC binding affinity records across 153 distinct MHCI molecules. These include molecules from various organisms: 7 BoLA, 1 Gogo, 7 H-2, 104 HLA, 19 Mamu, 11 Patr, and 4 SLA. For each allele, 25 random natural peptides of lengths 8, 9, 10, and 11 were generated as artificial negatives, totaling 100 artificial negatives per allele and 15,300 in the dataset ^20^. These artificial negatives were used in the training but excluded from evaluations in the five-fold cross-validation process. In line with previous research, experimentally determined IC50-binding values were transformed into binding affinity values on a [0,1] scale using the formula: 1 − log (*IC*_50_*nM*) / log (50000). The number of peptides per allele and the corresponding number of binders are detailed in Supplementary Table S1.

#### Binary_2024_

Binary_2024_ is a collection of automated benchmark datasets compiled from the Immune Epitope Database (IEDB) to evaluate various methods for predicting MHCI binding ^21^. Data measured by binary type from 2023 to 2024 were selected weekly to compile this independent dataset, removing records that overlap with BD2017. Especially, we removed the records that overlap with BD2017. The dataset contains 385 binding samples and 228 non-binding samples for 11 types of MHCI molecules with 501 peptides.

#### IEDB2016

IEDB2016 consists of 134,281 data records of MHC-peptide binding affinity, including 80 kinds of MHCII molecules, collected from IEDB ^22^ as of 2016. Similarly to the processing of BD2017, the affinity values in IEDB2016 were converted by the same transformation.

### PROBLEM FORMULATION

Given their amino acid sequences, this task involves predicting the binding affinity between MHC molecules and peptides, which is a regression problem. The binding affinity is primarily determined by the amino acids bound to the MHCI molecule’s binding groove, known as anchor residues. MHC molecules’ binding grooves can also vary depending on the peptide lengths. Here, we represent the MHCI pseudo-sequence as *Q* = (*q*_1_, *q*_2_, …, *q*_*L*_) the peptide sequence as *P* = (*p*_1_, *p*_2_, …, *p<SUB>L</SUB>′*), where each *p*_*i*_ and *q*_*i*_ denotes one of the 20 amino acids. In practice, we utilize a 34-length pseudo-sequence derived from source protein sequence to represent MHCI molecules ^23^. Given that most peptide sequences are shorter than 11 amino acids, we pad or truncate them to a length of 11 to ensure consistency in input dimensions.

### THE MODEL ARCHITECTURE

The Overview and the detailed architecture of DPCMHC are illustrated in Figure 1, which consists of an embedding module, an embedding module, a dual-padding convolutional module, a BiLSTM module, and an output module.

**Figure 1.**
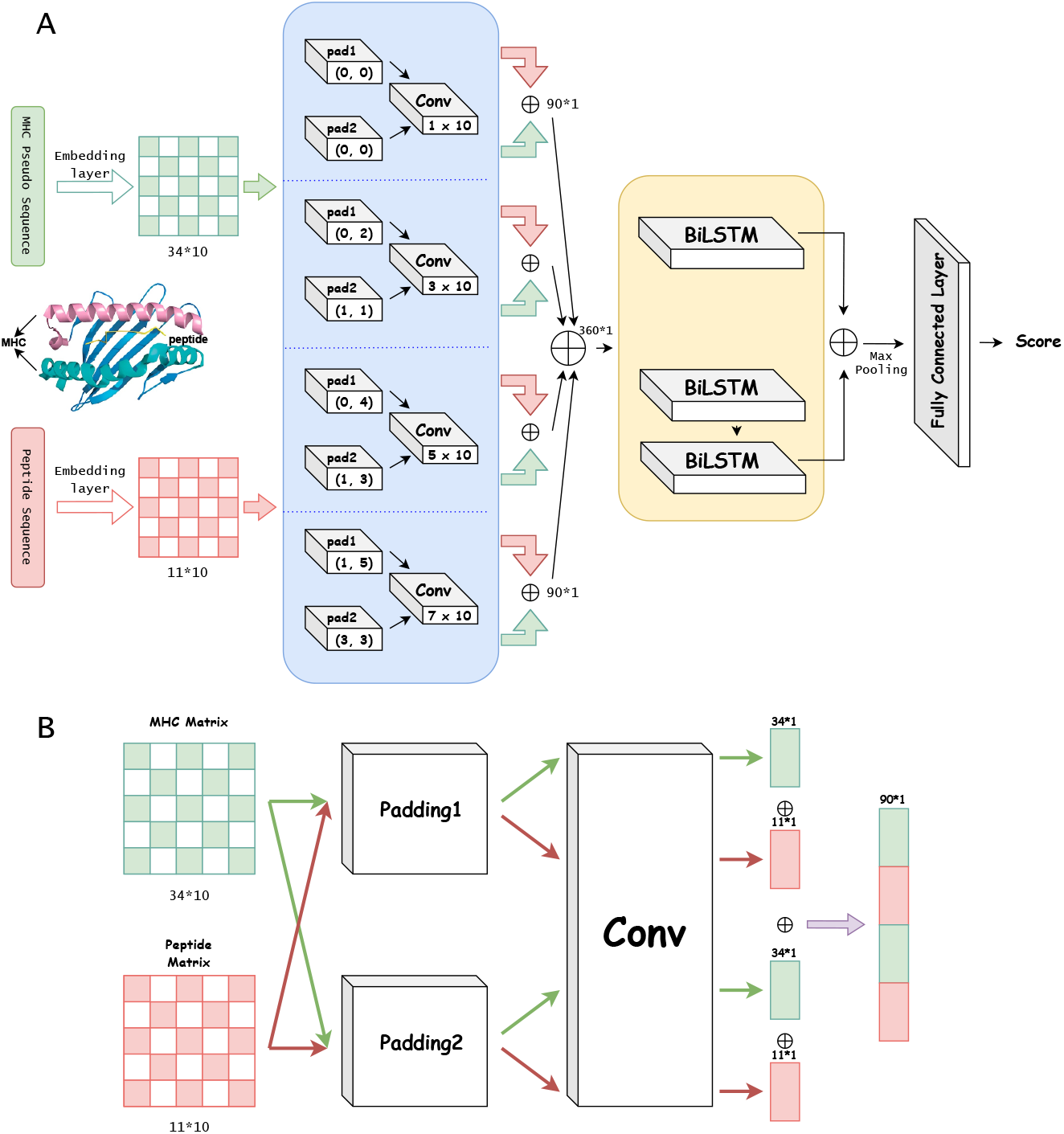
The model architecture of DPCMHC.(A) The overview of the model includes the four modules. (B) The details of the dual-padding module.

#### EMBEDDING MODULE

The MHCI pseudo-sequence and peptide sequence are represented by amino acid symbols, which can be viewed as linear chains of amino acids. We used a modified one-hot encoding method to convert these 20 standard amino acid characters into numerical values for calculations. Each amino acid is represented by a vector containing three ‘1’ and three ‘0’ ^24^. In addition, to enhance the distinction between different amino acids, we considered their polarity and charge status, classifying them into the four categories and using four bits (two ‘0’s and two ‘1’s) to represent each category: non-polar [0, 0, 1, 1], polar uncharged [0, 1, 0, 1], polar positively charged [0, 1, 1, 0], and polar negatively charged [1, 0, 0, 1]. Therefore, each amino acid is represented by a ten-element vector. Unknown residues *X* are represented by [1, 1, 1, 1, 0, 0, 0, 0, 0, 0]. This encoding results in a 34 *×* 10 matrix for each MHCI pseudo-sequence and an 11 *×* 10 matrix for each peptide sequence.

#### DUAL-PADDING CONVOLUTIONAL MODULE

This module comprises two components: dual-padding and convolution layers ^25^. We utilize convolutional kernels corresponding to the amino acids encoding length to effectively capture the sequential and structural information embedded in these representations. Unlike the square kernels typically used in image processing, our kernels are specifically designed to process the entire 10-bit representation of each amino acid in a single convolution step. This approach ensures that each convolution operation captures the full feature set of an individual amino acid while preserving the features of consecutive amino acids. This design enhances the model’s ability to learn relevant biological patterns. The kernels we use, sized 1 *×* 10, 3 *×* 10, 5 *×* 10, and 7 *×* 10, enable a more comprehensive integration of sequential data points, which is crucial for accurate prediction and analysis in this task.

Despite the input sizes for each encoded peptide and MHCI sequence are consistent, we employ two distinct padding strategies to optimize convolution operations. The first strategy adds padding only at the end of the sequence, while the second uses asymmetric padding, applying different lengths at the beginning and end. While both strategies use the same kernel size for each convolutional operation, their receptive fields differ except for *K*_0_. The substantial overlap between the receptive fields of both branches helps capture continuous information within the sequence, improving the model’s feature extraction capabilities compared to using a single padding strategy.

Specifically, For each convolution kernel *K*_*i*_ (1 ≤ *i* ≤ 4), the input matrices for MHC and peptides undergo dual-padding. The padding lengths at the beginning and end are denoted as *b*_*ij*_ and *e*_*ij*_ (1 ≤ *j* ≤ 2). This results in an MHC matrix of dimensions (*b*_*ij*_ + 34 + *e*_*ij*_) *×* 10 and a peptide matrix of dimensions (*b*_*ij*_ + 11 + *e*_*ij*_) *×* 10.

##### First padding way (*j* = 1)

Zero-padding is applied at the ends for all kernels except *K*_4_, i.e., *b*_*i*1_ = 0.

##### Second padding way (*j* = 2)

Padding is applied at both the beginning and the end, with *b*_*i*2_ ≠ 0 and *e*_*i*2_ ≠ 0.

For kernel *K*_0_, with a stride of 1 for each convolution kernel, the padding parameters are *b*_11_ = *b*_12_ = *e*_11_ = *e*_12_ = 0.

Table 1 below provides the specific values of *b* and *e*.

**Table 1.**
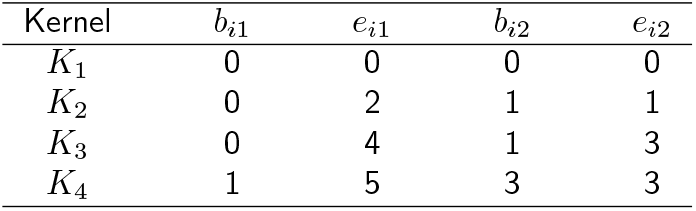
Values of different padding length *b* and *e* for 4 kernels.

| Kernel | $b_{i1}$ | $e_{i1}$ | $b_{i2}$ | $e_{i2}$ |
| --- | --- | --- | --- | --- |
| $K_1$ | 0 | 0 | 0 | 0 |
| $K_2$ | 0 | 2 | 1 | 1 |
| $K_3$ | 0 | 4 | 1 | 3 |
| $K_4$ | 1 | 5 | 3 | 3 |

#### BILSTM MODULE

While convolutional neural networks (CNNs) are effective at analyzing local input patterns and their surrounding context, they often struggle to capture long-range dependencies in sequential data. Bidirectional long short-term memory (BiLSTM) networks ^26^ address this limitation by processing information in both forward and backward directions, unlike standard LSTM models, which only rely on historical context. This bidirectional approach makes BiLSTM more effective for sequential modeling tasks. ^27^ Given BiLSTM’s superior ability to handle sequential data ^28^, we integrated it into this module to further enhance feature learning from the sequence. We employed two BiLSTM blocks: one with a single BiLSTM layer and another with two layers, each containing 128 hidden units. Both blocks extract features from the output of the dual-padding convolutional module. To clarify the BiLSTM architecture, the LSTM cell consists of four key components: the input gate (*i*_*t*_) with trainable weight matrices *W*_*xi*_, *W*_*hi*_, *W*_*ci*_, and bias *b*_*i*_; the forget gate (*f*_*t*_) with weights *W*_*xf*_, *W*_*hf*_, *W*_*cf*_, and bias *b*_*f*_ ; and the output gate (*o*_*t*_) with weights *W*_*xo*_, *W*_*ho*_, *W*_*co*_, and bias *b*_*o*_. The current cell state (*c*_*t*_) is updated based on a weighted sum of the previous cell state and the new information generated by the cell ^29^.

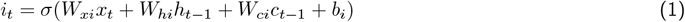

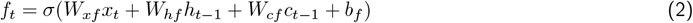

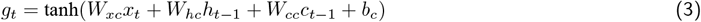

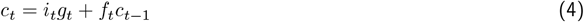

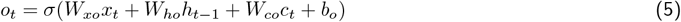

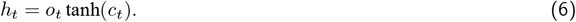

The outputs from these two BiLSTM blocks are then concatenated and pass through a max pooling layer, which pools the maximum features. This setup allows the model to capture both short-term and long-term dependencies in the sequence data, enhancing its ability to predict binding affinities accurately.

#### OUTPUT MODULE

This module consists of fully connected layers and an output layer. The fully connected layers contain 1024, 768, 640, and 384 neurons, each using ReLU as the activation function:

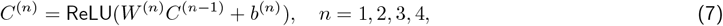

where *W* ^(*n*)^, *b*^(*n*)^ and *C*^(*n*−1)^ represent the weights, bias and output of the *n*-th fully connected layer, respectively. In addition, to avoid overfitting, we applied a dropout ^30^ after the first fully connected layer. The output layer contains a single neuron with a sigmoid activation function as follows, which is appropriate given that the binding affinity *ŷ* in datasets has been transformed between 0 and 1.

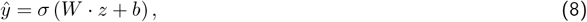

where *W* is the weight and *b* is the bias, respectively.

### MODEL TRAINING

As mentioned above, we treat this task as a regression problem of predicting the affinity between MHCI and peptides. Therefore, we use the mean squared error (MSE) loss function to train our model. The formula is as follows:

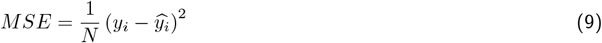

where *N* is the number of samples from training data, *y*_*i*_ refers to the experimentally measured binding affinity for the *i*th pair of MHCI and peptide, and 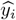 represents the predicted value of the same sample.

### EVALUATION METRICS

As in previous studies, for BD2017, only MHCI molecules with over 20 records and at least 3 binders were used for calculation. To classify peptides into binders and nonbinders, all peptides with an *IC*_50_ value < 500nM were classified as binders. However, unlike the area under the receiver operating characteristics curve (AUC) reported in DeepMHCI, we calculated the AUC for different lengths rather than averaging the AUC of each molecule. This approach better reflects the model’s generalizability and applicability in predicting the affinity of two given sequences. In addition, we employed the Pearson correlation coefficient (PCC) to evaluate the correlation between the predicted value and true values:

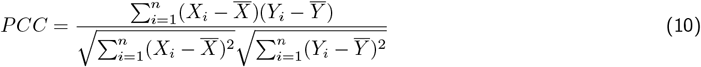

where *X*_*i*_ is the predicted value of DPCMHC and *Y*_*i*_ is the true binding affinity between MHC molecule and peptide for the same peptide length.

For Binary_2024_, since the dataset contains binary classification labels (0/1), we used AUC, Precision(PPV), Sensitivity, and F1-Score as evaluation metrics. Additionally, we used an ensemble of five models for validation. For IEDB2016, all the metrics were used to evaluate the performance of DPCMHC.

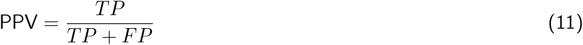

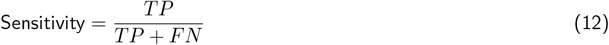

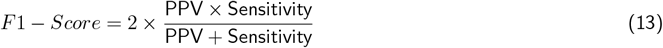

where TP, TN, FP, and FN represent true positive, true negative, false positive, and false negative, respectively.

## RESULTS

### COMPARISON OF DPCMHC AND BENCHMARK METHODS UNDER 5-FOLD CROSS-VALIDATION ON BD2017

We conducted a five-fold cross-validation on the BD2017 dataset to evaluate the performance of DPCMHC and compare it with DeepMHCI, DeepAttentionPan, and CapsNet-MHC under the same experimental settings. As shown in Table 2, DPCMHC outperforms all benchmark methods across multiple metrics for 9-mer, 10-mer, and 11-mer peptides. Notably, for 11-mer peptides, DPCMHC achieved the highest AUC of 0.881, surpassing DeepMHCI, DeepAttentinoPan, and CapsNet-MHC by 1.7%, 4.8%, and 5.0%, respectively. In addition, DPCMHC reached a PCC of 0.738 for 11-mer peptides, significantly higher than other models. The numbers in parentheses next to each peptide length in Table 2 represent their proportion in the overall dataset. The proportions of 8-mer and 12-mer peptides are quite small, at only 4.39% and 0.74%, respectively. While DPCMHC performs slightly worse than DeepMHCI for these two lengths, the difference is minor. However, for 9-mer, 10-mer, and 11-mer peptides, DPCMHC consistently outperforms the benchmarks, indicating its strong performance in most MHCI-peptide binding cases.

**Table 2.**
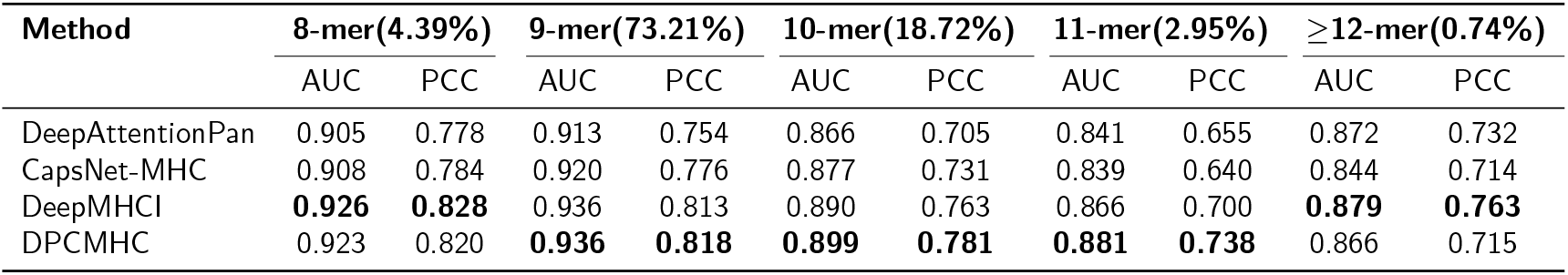
Performance of DPCMHC and benchmark methods on BD2017.

Figure 2 illustrates the AUC values achieved by the four methods for 10-mer and 11-mer peptides. The yellow bars represent the AUC of DPCMHC, while the purple, blue, and green bars correspond to DeepAttentionPan, CpaNet-MHC, and DeepMHCI, respectively. It is evident that the yellow bars are consistently higher than the others, particularly in the case of 11-mer peptides, clearly demonstrating the advantage of DPCMHC.

**Figure 2.**
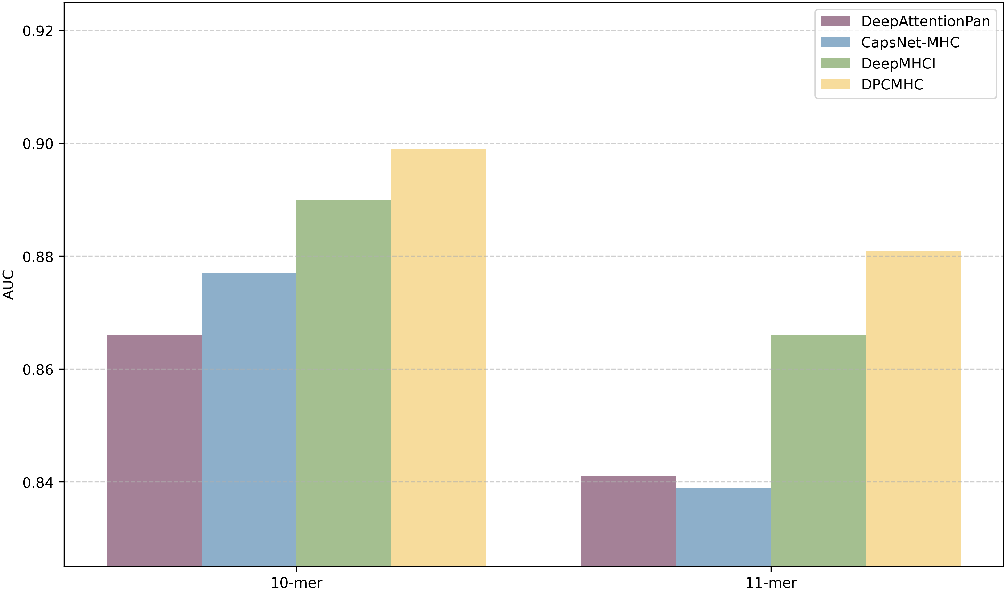
Performance comparisons of DPCMHC and benchmark methods were conducted for 10-mer, and 11-mer peptides using BD2017.

However, DPCMHC did not outperform the benchmark methods for 8-mer and ≥12-mer peptides. The reasons are as follows: 1) for 8-mer peptides, the input length is set to 11, followed by padding operations, which introduce excessive invalid information; 2) for ≥12-mer peptides, the input length setting leads to the omission of some amino acid information. Despite these challenges, DPCMHC’s performance remains close to the best-performing models, with differences of only 0.3% and 1.5% for 8-mer and ≥12-mer peptides, respectively.

### COMPARISON OF DPCMHC AND BENCHMARK METHODS ON Binary_2024_

We use Binary_2024_ as an independent test dataset to evaluate the performance of DPCMHC. Table 3 reports the performance of DPCMHC and benchmark methods on Binary_2024_, which consists of 406 records of 9-mer peptides and 207 records of 10-mer peptides. DPCMHC showed the best performance on the AUC, Sensitivity, and F1-Score for 9-mer peptides. For 10-mer peptides, DPCMHC achieved the highest Sensitivity and F1-Score. Specifically, for 10-mer peptides, DPCMHC achieved an F1-Score of 0.639, which was 5.8%, 25.8%, and 33.1% higher than those of DeepMHCI (0.604), CapsNet-MHC (0.508) and DeepAttentionPan (0.480), respectively. In addition, when evaluating all datasets, DPCMHC achieved the highest AUC (0.880), Sensitivity (0.738), and F1-Score (0.805).

**Table 3.** Performance of DPCMHC and benchmark methods on Binary_2024_.

| Metrics | DeepMHCI |  | DeepAttentionPan |  | CaspNet-MHC |  | DPCMHC |  |
| --- | --- | --- | --- | --- | --- | --- | --- | --- |
|  | 9-mer | 10-mer | 9-mer | 10-mer | 9-mer | 10-mer | 9-mer | 10-mer |
| AUC | 0.880 | **0.855** | 0.871 | 0.811 | 0.872 | 0.780 | **0.882** | 0.843 |
| Precision | 0.914 | **0.857** | **0.927** | 0.857 | 0.906 | 0.825 | 0.898 | 0.825 |
| Sensitivity | 0.797 | 0.467 | 0.736 | 0.333 | 0.783 | 0.367 | **0.803** | **0.522** |
| F1-Score | 0.847 | 0.604 | 0.820 | 0.480 | 0.840 | 0.508 | **0.848** | **0.639** |

In addition, we categorized the molecules in Binary_2024_ based on the number of binding peptides, resulting in two categories: 10-20 binders and 20-30 binders. Notably, DPCMHC significantly outperformed three benchmark methods. It is important to emphasize that each molecule had no more than 30 binding data points, underscoring DPCMHC’s robust performance with previously unseen MHCI molecules and limited data. The results are shown in Supplementary Figure S1. In Figure 3, the performance of DPCMHC is demonstrated using the H-2-Kb molecule. DPCMHC achieved a higher ROC ((*AUC*_*ROC*_ = 0.928)) compared to the other three methods, further validating its superior predictive capability and reliability in assessing peptide-MHC binding affinities. Additionally, DPCMHC outperformed the other methods in PRC (*AUC*_*PRC*_ = 0.93), indicating its greater accuracy in predicting binding outcomes and its consistent reliability across various levels of recall.

**Figure 3.**
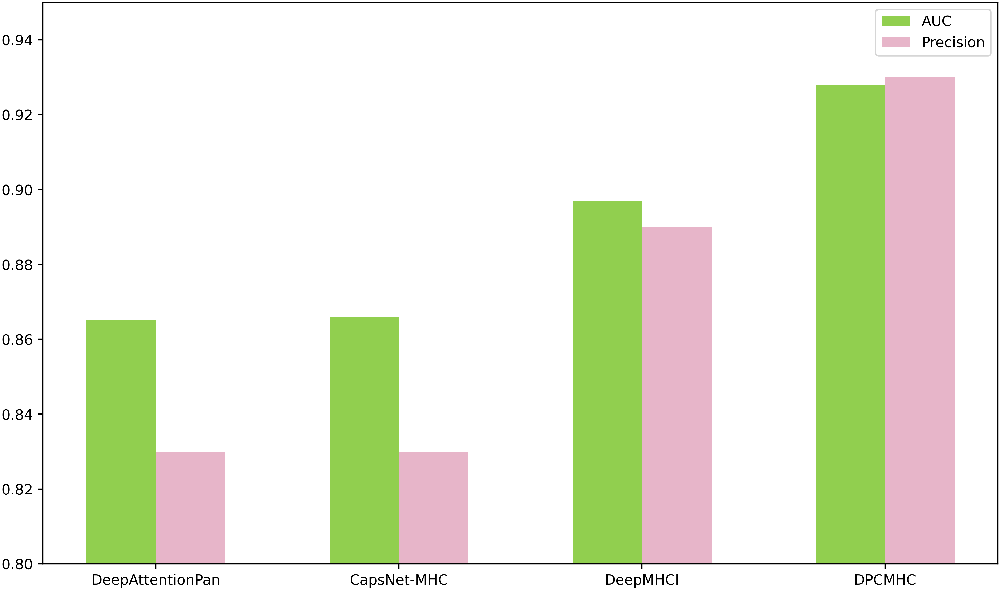
Performance comparison between DPCMHC and benchmark methods (DeepAttentionPan, CapsNet-MHC, and DeepMHCI) on H-2-Kb.

### COMPARISON OF DPCMHC AND BENCHMARK METHODS ON IEDB2016

To further assess the effectiveness and applicability of our model for predicting MHCII-peptide binding, we conducted a five-fold cross-validation on IEDB2016 dataset. We also compared the performance of DPCMHC with DeepAttentionPan and CapsNet-MHC under the same experimental conditions. Given the different peptide length distribution, we adjusted the input peptide length to 15. As shown in Table 4, DPCMHC outperformed the other methods on most metrics, achieving a PCC of 0.684, which is 13.6% and 8.6% higher than DeepAttentionPan and CapsNet-MHC, respectively. In addition, we calculate the average AUC and PCC of 61 MHC molecules, each with more than 40 records and at least 3 binders. Figure 4 illustrates that DPCMHC outperformed the other methods for over 85% of the molecules, demonstrating its robustness and exceptional generalizability across various MHC classes. Detailed AUC and PCC results are provided in Supplementary Table S2.

**Table 4.** Performance of DPCMHC and benchmark methods on IEDB2016.

| Metrics | DeepAttentionPan | CapsNet-MHC | DPCMHC |
| --- | --- | --- | --- |
| AUC | 0.795 | 0.809 | **0.837** |
| PCC | 0.602 | 0.630 | **0.684** |
| PPV | 0.720 | 0.699 | **0.743** |
| Sensitivity | 0.568 | **0.669** | 0.662 |
| F1-Score | 0.635 | 0.684 | **0.700** |

**Figure 4.**
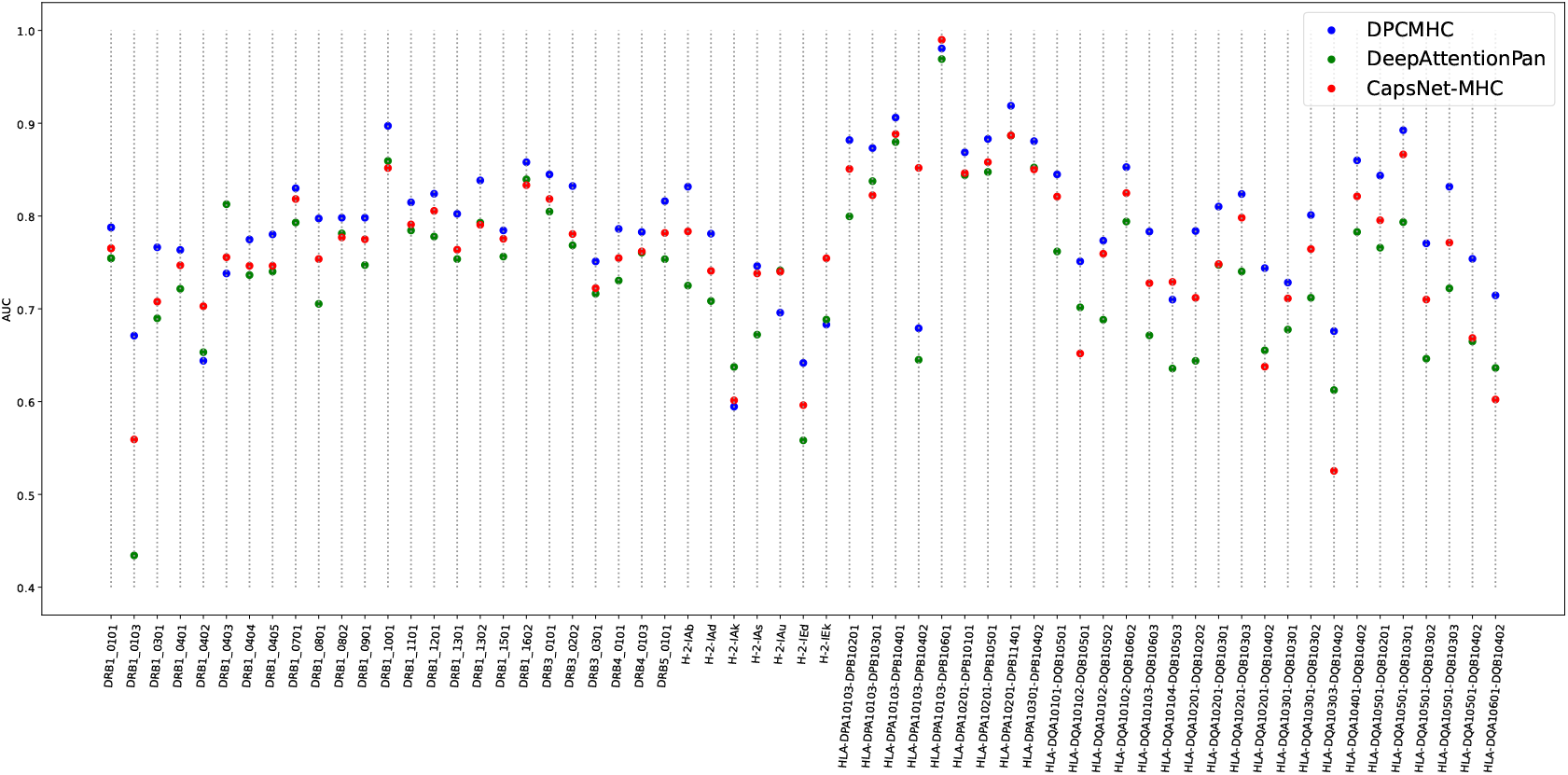
Comparison of DPCMHC (blue circles), DeepAttentionPan (green circles), and CapsNet-MHC (red circles) on 61 molecules of IEDB2016.

In addition, to further evaluate DPCMHC’s performance across different peptide lengths, we compared it directly with DeepMHCI on the IEDB2016 dataset, as DeepMHCI had outperformed DeepAttentionPan and CapsNet-MHC. Due to differences in peptide lengths between IEDB2016 and BD2017, we categorized the peptides into 3 categories: 9-12 mer, 13-19 mer, and ≥20 mer. To ensure reliable results, we only included lengths with more than 40 data points and at least three binders (as shown in Supplementary Table S2). We then calculated the AUC, PCC, and AUPRC for each category, with the results shown in Table 5. Unlike MHCI binding peptides, the most frequent peptide length in IEDB2016 is 13-19 mer. Although DPCMHC performs slightly worse than DeepMHCI in this range, it outperforms DeepMHCI in both the 9-12mer and ≥20 mer cases. Notably, for 9-12mer peptides, DPCMHC surpasses DeepMHCI in AUPRC by 9.28%, demonstrating its strength in shorter peptide predictions.

**Table 5.** Comparison of DPCMHC and DeepMHCI on different lengths in IEDB2016.

| Metrics | DeepMHCI |  |  | DPCMHC |  |  |
| --- | --- | --- | --- | --- | --- | --- |
| | 9-12 | 13-19 | $\geq 20$ | 9-12 | 13-19 | $\geq 20$ |
| AUC | 0.804 | **0.853** | 0.736 | **0.834** | 0.841 | **0.745** |
| PCC | 0.557 | **0.708** | **0.644** | **0.603** | 0.692 | 0.471 |
| AUPRC | 0.539 | **0.812** | 0.644 | **0.589** | 0.797 | **0.649** |

## RESULT ANALYSIS

### BINDING MOTIFS ANALYSIS

We used sequence logo ^31^ to visualize the binding motifs of MHCI molecules, highlighting the advantages of DPCMHC. The data, sourced from SwissProt ^32^, consists of 100,000 randomly selected 11-mer peptides. We calculated their binding scores and selected the top 1% (10,000 peptides) with the highest scores to generate the sequence logo. Using Seq2logo ^33^, the x-axis represents the alignment positions within the motif, with the total height of the letters (amino acids), indicating the amount of information content (or importance). The height of the individual letter corresponds to the frequency of specific amino acids at that position. Specifically, for Binary_2024_, we selected HLA-B08:01 and H-2-Kb as examples to illustrate this.

Coincidentally, the data points of HLA-B08:01 in Binary_2024_ are all 9-mer peptides. Figure 5A shows a scatter distribution of predicted values of DPCMHC (y-axis) and DeepMHCI (x-axis). As illustrated in Figure 5A, most green dots are above the diagonal line (y=x), indicating that DPCMHC provided higher predicted values for positive data points compared to DeepMHCI. Two samples, TLFMKEHNL and TLIMREHNL, with higher predicted values from DPCMHC, are highlighted. Notably, these peptides share identical amino acids at positions 2, 6, 7, 8, and 9. From the binding motifs generated by DPCMHC and DeepMHCI (Figure 5D and 5B, respectively), we observe that DPCMHC showed a strong preference for amino acid L at position 2. This explains why DPCMHC predicted higher values than DeepMHCI for these samples. Additionally, the main differences between the binding motifs from DPCMHC and DeepMHCI are mainly at positions 2 and 5, likely contributing to DPCMHC’s higher AUC compared to DeepMHCI and CapsNet-MHC on HLA-B08:01 in Binary_2024_.

**Figure 5.**
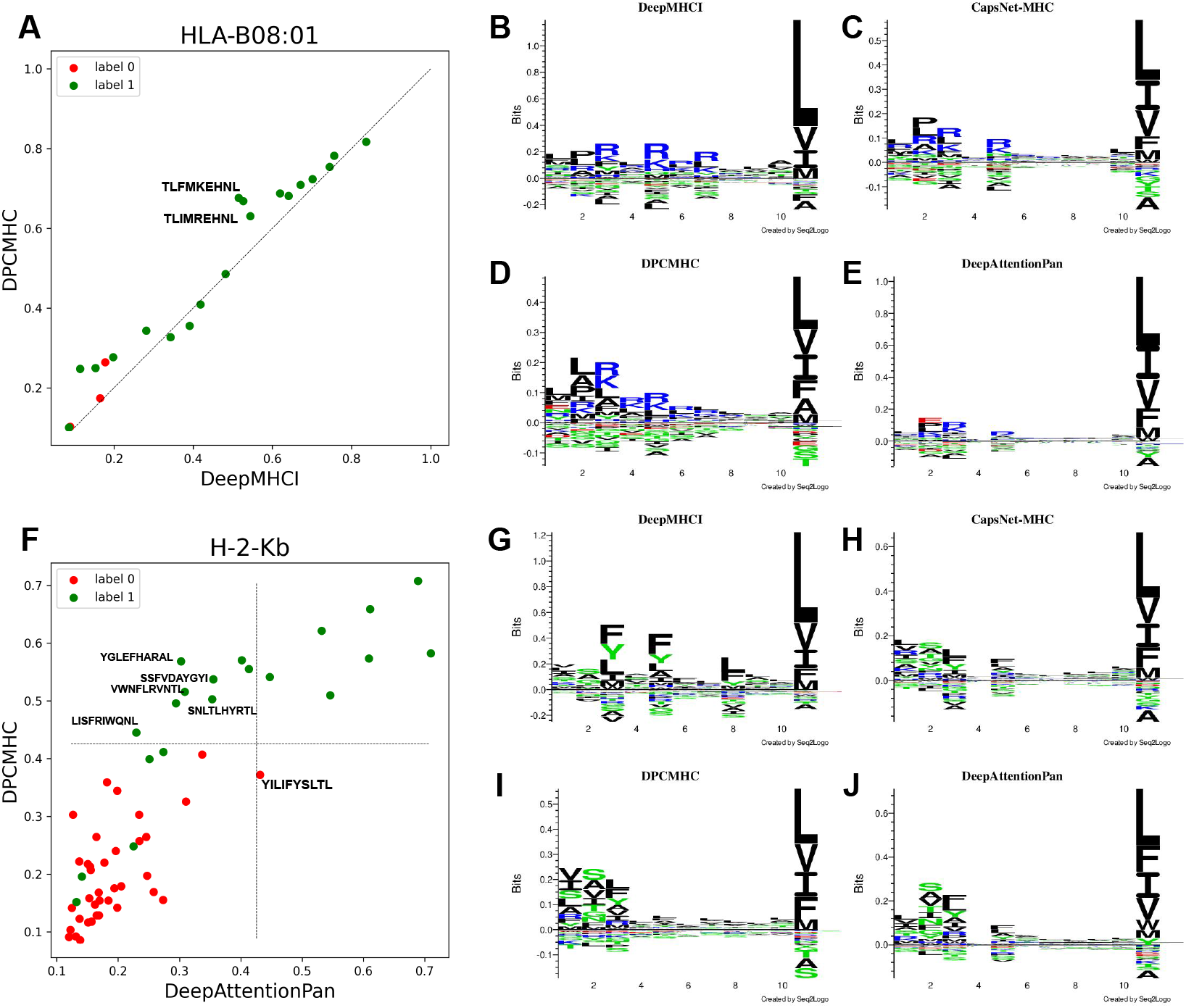
Comparison of binding motifs among DeepAttentionPan, CapsNet-MHC, DeepMHCI, and DPCMHC. (A) The scatter distribution of predicted scores for DPCMHC and DeepMHCI with the 9-mer of HLA-B08:01 in the Binary_2024_ dataset. (B-E) Binding motif of HLA-B08:01 under 9-mer peptides by DeepMHCI, CapsNet-MHC, DPCMHC, and DeepAttentionPan. (F) The scatter distribution of predicted scores for DPCMHC and DeepAttentionPan with the 10-mer of H-2-Kb in Binary_2024_ dataset. (G-J) The binding motif of H-2-Kb for 10-mer peptides by DeepMHCI, CapsNet-MHC, DPCMHC, and DeepAttentionPan.

From the binding motifs of DPCMHC and DeepMHCI (Figure 5D and 5B, respectively), both models show a preference for amino acids R and K at position 5. In addition, DPCMHC does not exhibit a strong preference for amino acids E, H, N, and L at positions 6, 7, 8, and 9 respectively. However, DPCMHC favors amino acid L at position 2, which explains why it predicted higher values than DeepMHCI for certain peptides. The primary difference between the two methods lies at position 2, which likely accounts for DPCMHC achieving a higher AUC than both DeepMHCI and CapsNet-MHC for HLA-B08:01 in Binary_2024_.

Moreover, the data points of H-2-Kb in Binary_2024_ are all 10-mer peptides. Figure 5F showed the scatter distribution of predicted values of DPCMHC (y-axis) and DeepAttentionPan (x-axis). As demonstrated, most green dots appeared in the upper area, indicating that DPCMHC successfully categorizes most positive and negative data points. However, we can see that DeepAttentionPan produced many false negative data points in the upper left area and a false positive data point. For the marked points in the left upper area, we found that amino acids S, V, and L appear at position 1. Comparing the binding motifs of DPCMHC and DeepAttentionPan (Figure 5I and 5J), DPCMHC shows a preference for amino acids V, S, and L, while DeepAttentionPan does not. This may explain the appearance of false negative points in the figure. In addition, focusing on position 3 of the motifs, it is evident that DPCMHC prefers amino acid L over F, while DeepAttentionPan shows the opposite preference. This may be why DeepAttentionPan predicted the peptide YGLEFHARAL as a false negative sample. For the false positive data point YILIFYSLTL, DeepAttentionPan has a preference for amino acid F at position 5, while DPCMHC does not share. This is consistent with the observation that DeepAttentionPan produces this false positive data point.

### ABLATION STUDY

In DPCMHC, we employed two padding ways before the convolutional operation to extract continuous sequence information. While existing methods typically pad sequence with zeros at the end, we aimed to validate the effectiveness of the dual-padding method by conducting five-fold cross-validation on the BD2017 dataset using only one padding way (referred to as DPCMHC-OP_*i*_). In addition, we evaluated the effectiveness of the BiLSTM module through five-fold cross-validation on BD2017. As shown in Table 6, DPCMHC exhibited the highest performance among the evaluated models. These results indicate that DPCMHC’s superior efficacy stems from both the dual padding methods and the incorporation of BiLSTM. The dual padding facilitates the extraction of continuous sequence information, while BiLSTM effectively captures the overarching dependencies inherent in sequential data.

**Table 6.**
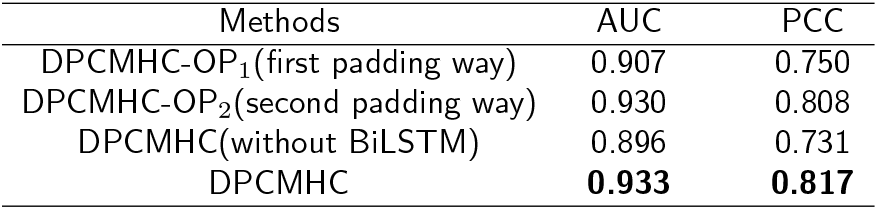
Performance between DPCMHC and DPCMHC-OP with BiLSTM overall data of BD2017.

| Methods | AUC | PCC |
| --- | --- | --- |
| DPCMHC-OP<sub>1</sub>(first padding way) | 0.907 | 0.750 |
| DPCMHC-OP<sub>2</sub>(second padding way) | 0.930 | 0.808 |
| DPCMHC(without BiLSTM) | 0.896 | 0.731 |
| DPCMHC | **0.933** | **0.817** |

## CONCLUSION

In this work, we developed DPCMHC, a new deep-learning method based on dual-padding convolution for predicting the affinity between MHCI molecules and peptides. Specifically, the embedded sequences were padded in two different ways and concatenated, helping to better extract continuous sequence information. Additionally, we used four convolutional kernels of different sizes to capture information from adjacent amino acids of varying sizes. Furthermore, to capture relationships between amino acids from both directions, we utilized BiLSTM, leveraging its bidirectional structure.

Through comprehensive experimental investigation, we found that DPCMHC performed better than three benchmark methods on 9-mer, 10-mer, and 11-mer peptides, and performed close to the best method under other circumstances. Moreover, DPCMHC demonstrated commendable generalization capabilities, as evidenced by its ability to predict the binding of MHCII and peptides. However, there are still areas for improvement. First, the input length and padding length currently limit performance on 8-mer and ≥12-mer peptides. Second, the number of neurons in BiLSTM and fully connected layers results in longer training time. Future work will focus on developing more effective embedding and padding strategies and adjusting the neurons to shorten the training time.

## DATA AND CODE AVAILABILITY

All source code and data are available at GitHub https://github.com/sourcescodes/DPCMHC.

## SUPPLEMENTAL INFORMATION INDEX

Figures S1. Performance of DPCMHC on some molecules of Binary_2024_

Table S1. A detailed description of dataset BD2017

Table S2. Supplementary for the performance on IEDB2016

