## Supplementary Material for "DPCMHC: efficient prediction of MHC-peptide binding affinity by deep learning based on dual-padding convolution"

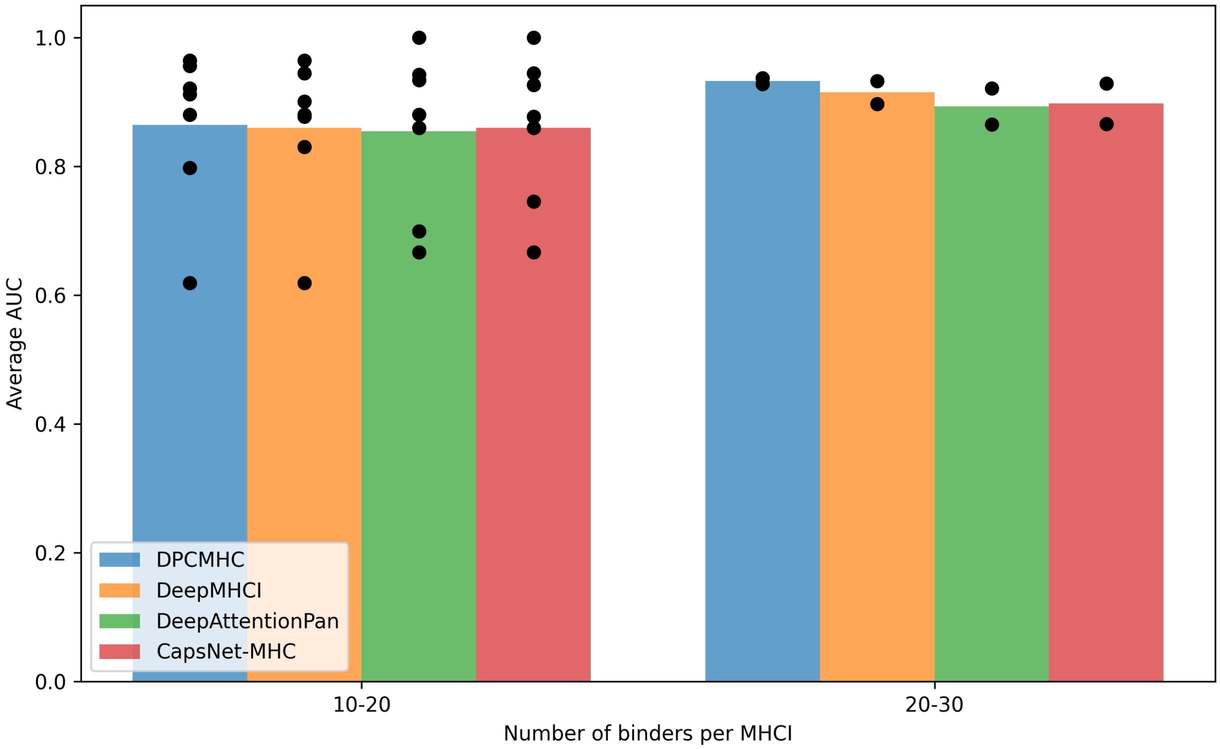


Figure S1: Performance comparison between DeepAttentionPan, DeepMHCI, CaspNet-MHC and DPCMHC on two categories of MHCI molecules in $\text{Binary}_{2024}$ with different number of binding peptides. Each bar shows the average performance of each method in terms of AUC on all MHC molecules in a certain category. Each scatter point represents the performance of an MHC molecule.

Table S1: Composition of BD2017

| Allele | MHCs | Binders | Peptides |
|---|---|---|---|
| BoLA | 7 | 610 | 1264 |
| Gogo | 1 | 6 | 15 |
| H-2 | 7 | 2716 | 9771 |
| HLA | 104 | 40841 | 156818 |
| Mamu | 19 | 4969 | 14018 |
| Patr | 11 | 1062 | 3710 |
| SLA | 4 | 201 | 389 |

Table S2: Performance of DPCMHC and competing methods on the 61 molecules of IEDB2016

| **Methods** | **AUC** | **PCC** |
|---|---|---|
| DeepAttentionPan | 0.738 | 0.479 |
| CapsNet-MHC | 0.763 | 0.523 |
| DPCMHC | 0.793 | 0.581 |
